# A virally encoded high resolution screen of cytomegalovirus host dependencies

**DOI:** 10.1101/2023.10.30.564793

**Authors:** Yaara Finkel, Aharon Nachshon, Einav Aharon, Tamar Arazi, Elena Simonovsky, Martina Dobesova, Avi Gluck, Tal Fisher, Richard J. Stanton, Michal Schwartz, Noam Stern-Ginossar

## Abstract

Genetic screens have transformed our ability to interrogate cellular factor requirements in infection, yet current approaches are limited in their sensitivity, biased towards early stages of infection and provide only simplistic phenotypic information which is often based on infected cell survival. Here, by engineering human cytomegalovirus to express sgRNA libraries directly from the viral genome, we developed a sensitive, versatile, viral centric approach that allows profiling of different stages along viral infection in a pooled format. Using this approach, which we termed VECOS (Virus Encoded CRISPR-based direct readOut Screening system), we identified hundreds of novel host dependency and restriction factors and quantified their direct effects on viral genome replication, viral particle secretion and infectiousness of secreted particles, providing a multi-dimensional perspective on viral-host interactions. These high resolution measurements reveal that perturbations that alter late stages in HCMV life cycle mostly regulate HCMV particle quality rather than quantity, defining correct virion assembly as a critical stage that is heavily reliant on viral-host interactions. Overall, VECOS facilitates systematic high resolution dissection of human proteins’ role along the infection cycle, providing a roadmap for in-depth dissection of host–herpesvirus interactions.

## Main

Understanding how viral and host factors interact and how perturbations impact infection is the basis for designing efficient antiviral interventions and for deciphering the molecular processes that take place in infected cells. CRISPR screens enable unbiased interrogation of gene function in diverse biological processes. However in the context of infection, insights are still mostly confined to early steps in the viral cycle, predominantly entry ^1^, greatly limiting the breadth of biology that could be uncovered. In a typical infection-related screen, Cas9 and a single guide RNA (sgRNA) are introduced into cells, the gene-edited cells are then challenged with a virus and the cells compete based on the fitness effect of the genetic perturbations. A major limitation of this screening approach is that it probes the cell state (survival) and not the direct effects of perturbation on the virus. Furthermore, these screens provide only simplistic phenotypes such as viral dependency or restriction, conflating genes that act via different mechanisms and making it impossible to identify proteins and processes that have more complex pleiotropic effects. Thus, extensive follow-up studies are needed to disentangle relevant hits, and our ability to effectively probe viral-host interfaces remains a great challenge.

To overcome these limitations, and to generate a platform that can transform the way we dissect host dependencies, we set-out to establish a screening system that directly measures the effect of genetic perturbations on viral propagation. For this we envisioned a system in which a sgRNA library is expressed directly from the viral genome. In such an approach, after a virus encoding an sgRNA infects a cell that expresses Cas9, the respective gene will be knocked-out in this cell. In cells where the targeted cellular protein affects the propagation of the virus, the level of sgRNAs (which are part of the viral genome) serve as a direct readout for the effect on virus propagation; sgRNAs that target genes that are essential or restrict the virus are expected to be under-represented or enriched, respectively (Figure 1a). Since thousands of viral genome copies are synthesized in each replication cycle, the signal is naturally amplified, making this screening approach extremely sensitive, potentially facilitating the detection of even small positive and negative effects on viral propagation. Furthemore, since sgRNA abundance directly reports on the virus levels, quantifying the sgRNAs at different compartments or stages of infection can inform on the stage in infection that is being affected by each perturbation, enabling high resolution analysis of cellular proteins’ effects on viral propagation.

**Figure 1.**
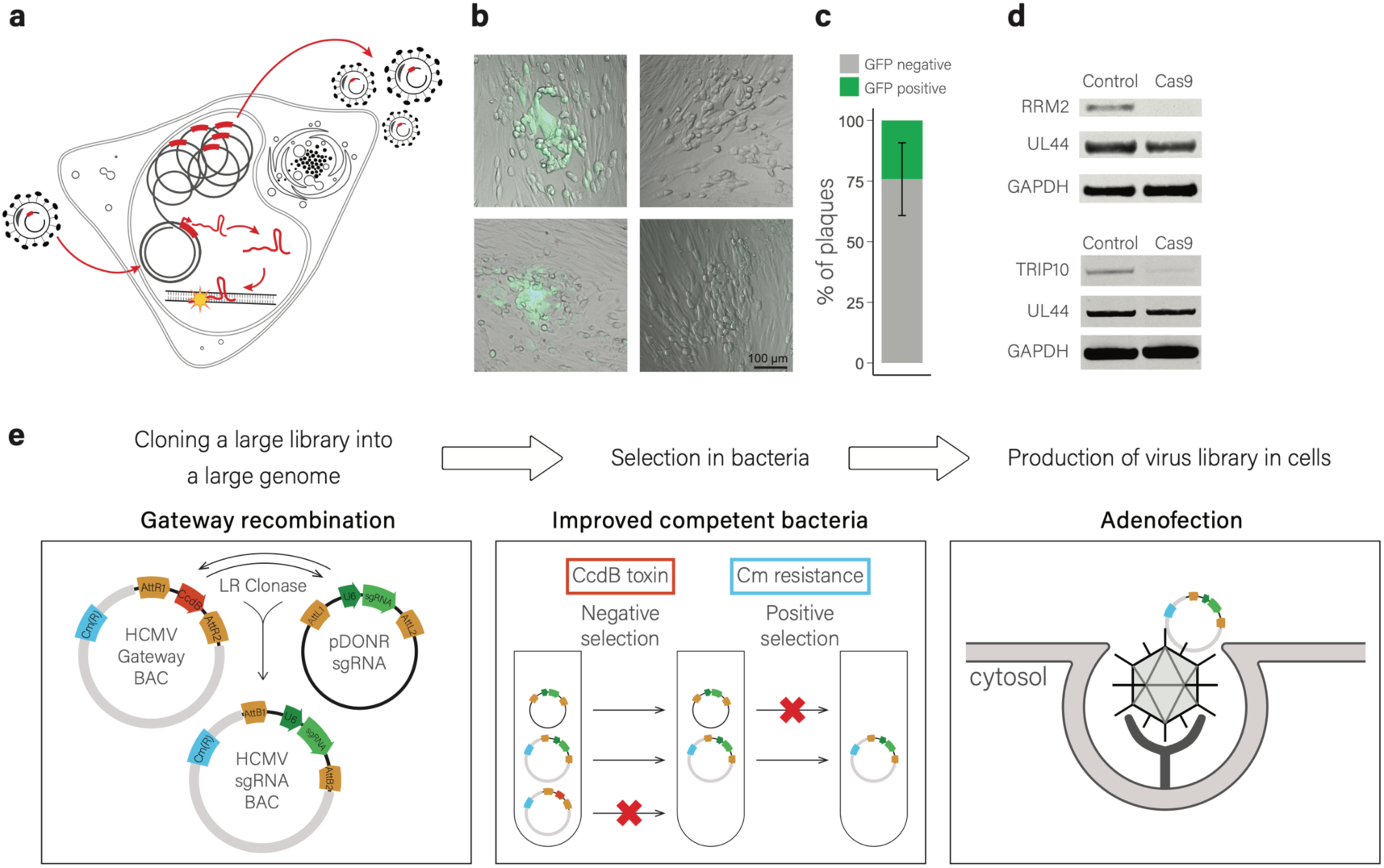
Establishment of VECOS strategy to express sgRNAs directly from the HCMV genome. a. The VECOS platform. A virus encoding an sgRNA in its genome infects a cell expressing Cas9. Upon infection, the sgRNA is expressed and interacts with Cas9 to induce site-specific mutations in the host genome. The viral genome replicates in the cells and produces new progeny containing the encoded sgRNA sequence. The level of viral genomes depends on the effect of the mutation induced by the virally encoded sgRNA. b. Examples of HCMV plaques in infected fibroblasts that are GFP positive (left) or GFP negative (right). c. Percentages of GFP positive and negative plaques formed by viruses budding from Cas9- expressing fibroblasts infected with HCMV encoding GFP and a sgRNA targeting GFP. Error bars represent the standard deviation of three biological replicates. d. Immunoblot analysis of RRM2 (top) and TRIP10 (bottom) protein expression at 3 dpi. Proteins were extracted from Cas9- or mCherry- (control) expressing fibroblasts infected with HCMV expressing either RRM2 or TRIP10 sgRNAs. GAPDH and UL44 were used as host and viral loading controls, respectively. e. Main steps in generating a VECOS library. Step I: A Custom Donor plasmid (pDONR-sgRNA) containing an sgRNA expression cassette (green) flanked by recombination sites (orange), and an HCMV BAC (HCMV-GW) containing the CcdB toxin gene (red) flanked by recombination sites (orange) and Chloramphenicol resistance gene (blue) were designed to facilitate introduction of a sgRNA library into the HCMV BAC via Gateway recombination. Step II: The Gateway recombination reaction products were transformed into optimized electrocompetent bacteria, CcdB toxin was used for negative selection (eliminating the parental BAC) and Chloramphenicol (Cm) was used for positive selection. Step III: The BAC library was introduced into fibroblasts via Adenofection to produce an infectious HCMV virus encoding the sgRNA.

As a proof of concept for the feasibility of such an approach we focused our efforts on the β-herpesvirus human cytomegalovirus (HCMV), a dsDNA virus, which is a pervasive pathogen that encodes for hundreds of proteins and has a slow and intricate infection cycle. HCMV genome engineering has been greatly enhanced by the cloning of the full HCMV genomes into bacterial artificial chromosomes (BACs)^2^. However, due to the large size of BACs, genomic manipulations are inefficient, making it challenging to generate complex libraries. Therefore to clone sgRNA libraries into the HCMV genome, we utilized an *in-vitro* Gateway reaction ^3^. We generated an HCMV BAC which also contains a GFP reporter with the Gateway acceptor site (HCMV-GW) and in parallel, we constructed a donor plasmid, with a Gateway cassette flanking the U6 promoter and an sgRNA scaffold (pDONR-sgRNA). Combining the HCMV-GW and pDONR-sgRNA in a Gateway reaction results in the insertion of an sgRNA under the U6 promoter into the HCMV genome (Supplementary figure 1a).

To test if sgRNAs are functional when expressed directly from HCMV, we cloned several individual sgRNAs; an sgRNA that targets GFP and two sgRNAs targeting human genes, TRIP10 and RRM2. We then created HCMV viruses by transfecting these HCMV BACs into fibroblasts and collecting the viruses, each of which contains a different sgRNA. Infection with the virus that contains an sgRNA targeting GFP, in cells that express Cas9, provided a fast way to assess targeting as the virus also encodes for GFP. Indeed, more than 70% of the viruses that were collected from Cas9-expressing fibroblasts (Cas9-cells), lost their GFP expression (Figure 1b and 1c), illustrating that sgRNAs expressed from the virus could generate functional Cas9 complexes. We next analyzed if cellular genes could also be efficiently targeted. Viruses containing sgRNAs targeting *TRIP10* or *RRM2* were used to infect Cas9-cells, and the host DNA was extracted at different time-points post infection. We then measured the proportion of mutations at the targeted loci along a time course of infection by deep sequencing. We found that indels start accumulating in these loci as early as 10 hpi, and continue to increase with infection progression (Supplementary Figure 1b and 1c). The kinetics of mutation accumulation are consistent with previous studies, where both Cas9 and the sgRNA were expressed from the cellular genome^4^. In addition, we measured the protein expression of TRIP10 and RRM2 in Cas9 and mCherry (control)-expressing fibroblasts infected with viruses expressing sgRNA targeting *RRM2* or *TRIP10*. At 3 dpi we observed significant reduction in RRM2 and TRIP10 expression in Cas9-cells relative to the control cells (Figure 1d), further illustrating sgRNAs expressed from HCMV can target cellular proteins. Taken together, these results illustrate the feasibility of expressing sgRNAs directly from HCMV and we named this targeting approach VECOS (Virus Encoded CRISPR-based direct readOut Screening system).

We next designed an sgRNA library that targets cellular genes. Since in VECOS targeting occurs after cells are infected we focused on 2,240 cellular genes that are upregulated in HCMV infection ^5^. For each of these genes we designed four sgRNAs and to these we added 500 non-targeting sgRNAs, adding up to a total of 9,460 sgRNAs (Supplementary table 1). The sgRNAs were cloned into the HCMV-GW using a Gateway reaction (Figure 1e, left panel). We then used negative and positive selection to specifically amplify bacteria that contain BACs with sgRNAs (Figure 1e, middle panel) and extracted BACs that retained 89% of our original library. To create HCMV viruses, BACs are transfected into fibroblasts to recover functional viruses. To increase the efficiency of this step we used inactivated adenovirus particles as carriers for the BAC DNA ^6–9^ (Figure 1e, right panel). Although it is likely there is still room for optimization, we nevertheless generated a virus stock that represented 78% of our original library (Supplementary figure 1d and supplementary table 1), illustrating it is possible to create highly complex HCMV libraries.

We next used the VECOS host targeting library to examine the effects of cellular proteins on HCMV propagation. We infected Cas9-cells in triplicates, using low multiplicity of infection (MOI=0.05) to minimize the chance of expressing multiple sgRNAs in the same cell. Since the sgRNAs are encoded in the viral genome, VECOS permits several rounds of selection, boosting the selection signal. We therefore performed three sequential passages in Cas9-cells. In each passage we collected the supernatant containing virus at 10 dpi and we also harvested the infected Cas9-cells (which contain viral DNA) at the time of virus collection. We then used a portion of the collected supernatant to perform another round of selection by again infecting fresh Cas9-cells at MOI of 0.05 (Figure 2a). At the end of the experiment, the sgRNA abundance was measured from the infected Cas9-cells and from the infectious virus that was collected, respectively. The distribution of sgRNA in infectious viruses was measured by infecting WT fibroblasts with the collected supernatant and analyzing sgRNA distribution in these cells at 3 dpi. Since these cells do not express Cas9, there is no selection along this infection and the sgRNAs serve solely as barcodes, where their relative abundance reflects the sgRNAs distribution in the infectious particles in the supernatant collected (Supplementary figure 1e).

**Figure 2.**
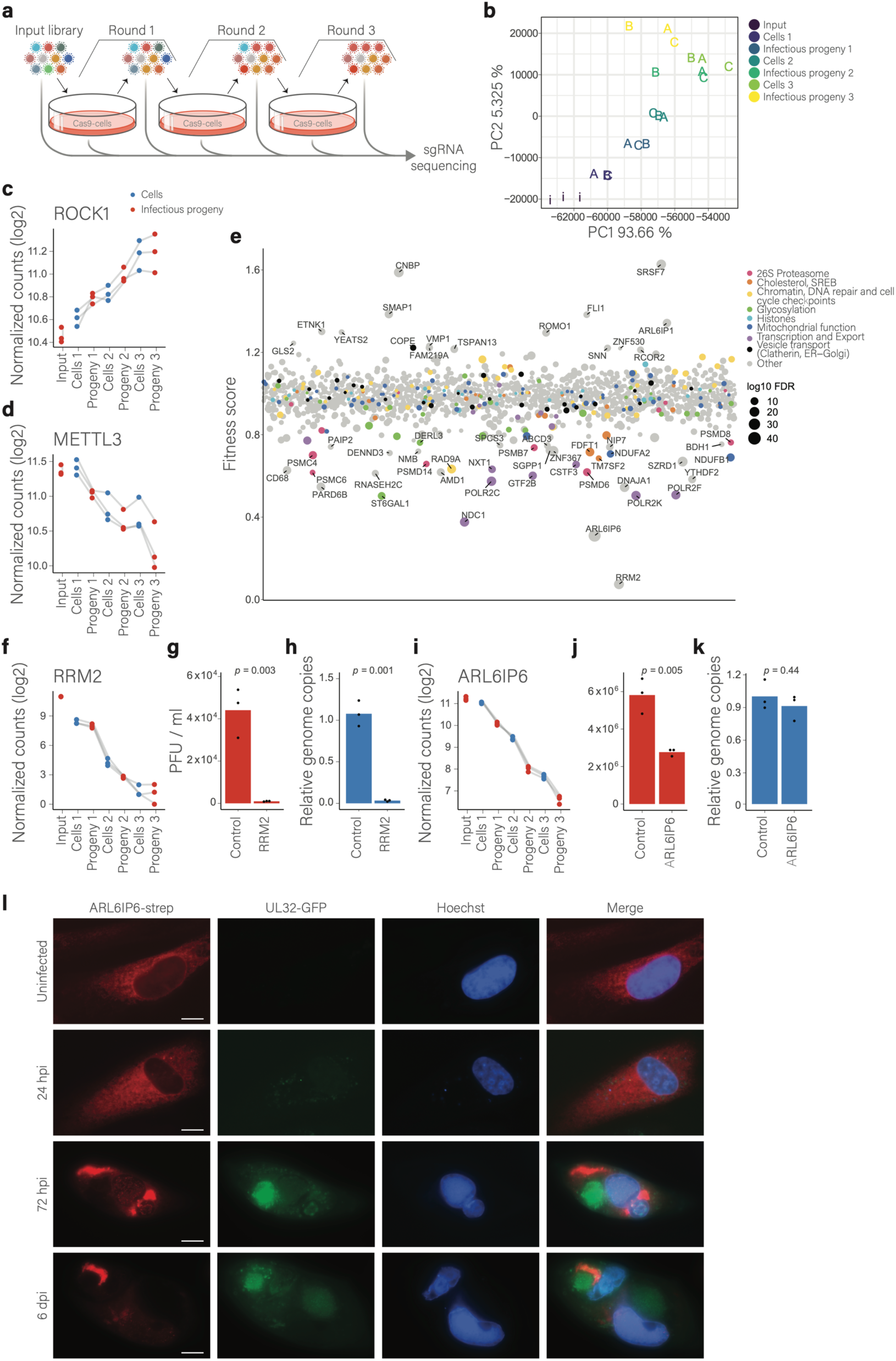
VECOS based host screen identifies host factors that are required or limit HCMV propagation. a. Host targeting VECOS screen. Cas9-expressing fibroblasts (Cas9-cells) were infected with the HCMV sgRNA library (MOI=0.05). At 10 dpi cells and supernatant were collected and sgRNA abundance in the cells and in the infectious virus in the supernatant was measured by deep sequencing. Progeny viruses were used to perform two additional rounds of selection in cas9-cells. The screen was performed in three independent biological replicates. b. Principal component analysis (PCA) of relative sgRNA abundance of the top 1% most varying sgRNAs. Percentages represent the explained variance. Colors indicate the stage of screen for each sample, and the data points are indicated as letters representing the replicates (A, B, C) and the input library (i). c and d. Relative levels of sgRNAs (geometric mean) targeting the indicated genes, ROCK1 (c) and METTL3 (d) along the screen (cells, blue; infectious progeny, red). Gray lines connect the values of the same biological replicate. e. The fitness score of genes targeted in the screen. Point size reflects the significance (-log10 FDR) and genes belonging to groups of interest are highlighted in color. The X axis is random. f. Relative levels of sgRNA (geometric mean) targeting RRM2 along the screen (cells, blue; infectious progeny, red). Gray lines connect the values of the same biological replicate. g. HCMV titers in control and RRM2 knockout cells. Viral supernatants were collected at 7 dpi from RRM2 knockout cells and control cells and transferred to wild-type fibroblasts. Infected cells were analyzed by flow cytometry 48 hpi. Plaque forming units (PFU) per ml were calculated from the percentages of GFP positive cells and the infection volume. h. Viral genome replication in control and RRM2 knockout cells. DNA from RRM2 knockout cells and control cells infected with HCMV was extracted at 3 dpi and the levels of host and viral DNA in each sample was measured by quantitative real-time PCR (qPCR). Viral DNA levels were normalized to host DNA and to the control samples mean. i. Relative levels of sgRNAs (geometric mean) targeting ARL6IP6 at different stages along the screen (cells, blue; infectious progeny, orange). The geometric means of sgRNAs targeting the gene are shown. Gray lines connect the values of the same biological replicate. j. HCMV titers in control and ARL6IP6 knockout cells. Viral supernatants were collected at 7 dpi from ARL6IP6 knockout cells and control cells and transferred to wild-type fibroblasts. Infected cells were analyzed by flow cytometry 48 hpi. PFU per ml were calculated from the percentages of GFP negative cells and the infection volume. k. Viral genome replication in control and ARL6IP6 knockout cells. DNA from ARL6IP6 knock-out cells and control cells infected with HCMV was extracted at 3 dpi and the levels of host and viral DNA in each sample was measured by qPCR. Viral DNA levels were normalized to host DNA and to the control samples mean. l. Fluorescence microscopy of fibroblasts expressing a strep-tagged version of ARL6IP6 (red) either uninfected or infected with HCMV strain expressing GFP fused to the tegument protein UL32, which localizes to the viral assembly compartment, (green) at 24, 72 hpi and 6dpi. Nuclei are stained with Hoechst (blue). Scale bar is 10 μm. In all plots, p-values were calculated using Student’s t-test. Bars represent the means of biological replicates.

Overall, correlation was high between all pairs of samples (Pearson’s R > 0.91), and no dropout of sgRNA was observed (Supplementary figure 2a) illustrating the complexity of the library was largely preserved in all passages. The abundance of the vast majority of non-targeting sgRNA did not dramatically change along the selection rounds (Supplementary figure 2b). Furthermore, PCA analysis performed on the sgRNA abundance values of most varying sgRNAs revealed a pattern compatible with consistent and directional change along rounds of selection (Figure 2b).

To identify genes whose targeting significantly affects HCMV propagation, we initially used MAGeCK ^10^, a widely used algorithm that allows the identification of significant hits in CRISPR/Cas9 knockout screens. The sgRNA distribution in the infectious progeny that budded out from Cas9-cells in each selection round was compared to the input library. The number of significantly changing genes increased with every passage, reaching 59 at the last round of selection (Supplementary figure 2c, FDR < 0.05). This shows that VECOS produces a robust gene-level selection signal that is strengthened along additional rounds of selection. Furthermore, we captured genes that we have previously reported to act as restriction factors (ROCK1) ^11^ or dependency factors (METTL3) ^12,13^ supporting the validity of the identified hits (Figure 2c and 2d).

MAGeCK analysis allows comparison of sgRNA distribution between two conditions. However, our screen provides longitudinal changes along selection rounds and this information is not incorporated into the analysis. We therefore devised a mixed-model regression approach that assesses the change in sgRNA distribution along the passages in both the cells and the viruses (see Methods), assigning for each gene a fitness score (reflecting the effect size, which is the fold-change in sgRNA abundance in one round of selection) and a p-value (reflecting the significance of the linear relationship between selection round and sgRNA abundance). Hits were selected by filtering on statistical significance and minimum effect size (both thresholds were calculated based on random sampling of the non-targeting sgRNAs). We further required that at least two sgRNAs will have a statistically significant linear change. Overall this analysis identified 258 genes that significantly affected HCMV propagation, 101 restriction factors and 157 dependency factors (Figure 2e, supplementary figure 2d and supplementary table 2, https://finkely.shinyapps.io/vecos_data_viewer_2023). Reassuringly, all but three of the MAGeCK hits were identified in our regression-based analysis. These three failed to be included in the regression results due to insufficient consistency between sgRNAs (Supplementary figure 2e).

Pathway enrichment analysis reveals that dependency factors are enriched in several functional categories such as glycosylation (p=3.4 × 10^−4^, hypergeometric test), possibly indicating a requirement for proper glycosylation of HCMV glycoproteins ^14^, RNA polymerase II components (p=3.6 × 10^−7^, hypergeometric test), as well as 26S proteasome subunits (p=7 × 10^−7^, hypergeometric test). A possible confounding factor in the screen is our inability to separate between effects on cell viability, which will indirectly harm viral propagation, and direct effects on virus propagation. To address this issue, we compared the fitness score of each gene in our screen to a viability score measured by quantifying the effects of gene targeting on primary fibroblasts growth following 10 days in cell culture ^15^. There was no significant correlation between the effects on viral propagation and the effects on cell viability in uninfected (Pearson’s R=0.06, supplementary figure 2f) or HCMV infected fibroblasts (Pearson’s R=0.01, supplementary figure 2g), indicating most of the signal we capture does not originate from indirect effects on cell viability. The only exceptions were proteasome related genes, whose deletion reduced fibroblasts viability and HCMV propagation, raising the possibility that the effect of proteasome depletion on viral replication may stem from a negative effect on cell viability (Supplementary figure 2f). However, this does not rule out specific roles for the proteasome in HCMV infection, as reflected by previous reports ^16^ and also described below. Notably, host factors identified are significantly enriched with proteins that directly interact with HCMV proteins (p=0.004, hypergeometric test) ^17^, providing additional independent support for their roles in virus propagation.

By far the strongest hit in the screen (Fitness score=0.07, with some of its sgRNAs completely disappearing from the population, figure 2f) was ribonucleotide reductase regulatory subunit M2 (RRM2), which catalyzes the rate-limiting step for production of dNTPs. CRISPR knockout of RRM2 severely hampered HCMV propagation (Figure 2g) and viral DNA replication (Figure 2h). Indeed RRM2 inhibitors were shown to inhibit HCMV propagation ^18^. The potent effects of RRM2 depletion and the approved clinical use of RRM2 inhibitors ^19^, support the idea of combining RRM2 inhibitors with traditional anti HCMV therapy (such as GCV) to increase the anti-HCMV activity ^18^. The extent by which RRM2 affects HCMV replication, and its absence from the list of hits of survival-based CRISPR screens performed on CMV infection ^15,20^ highlight the unique strength of VECOS and its ability to identify a wide range of host factors.

Another strong dependency factor identified in the screen is ADP ribosylation factor like GTPase 6 Interacting Protein 6 (ARL6IP6, Figure 2i, fitness factor = 0.31). ARL6IP6 is an uncharacterized membrane protein that was shown to localize to the inner nuclear membrane. We confirmed that CRISPR knockout of ARL6IP6 severely reduced HCMV propagation (Figure 2j) but not via inhibition of viral DNA replication (Figure 2k). Furthermore imaging of ARL6IP6 in uninfected fibroblasts and along HCMV infection revealed that late in infection ARL6IP6 relocalizes to cytoplasmic structures, that partially engulf the assembly compartment (Figure 2l), reminiscent of ER-derived structures that were shown to form late in infection ^21^ Indeed, ARL6IP6 localizes to the same structures as the ER marker calnexin (Supplementary figure 2h).

Beyond the superb sensitivity of VECOS and its ability to capture post-entry events, it also facilitates systematic dissection of the stage in infection that is affected by each perturbation, providing immediate insights into the mechanism of action. Since the sgRNAs are embedded in the viral genome, the relative abundance profiles of sgRNAs in different compartments indicates the step in the viral life cycle that is affected by each perturbation. For example, the abundance profiles of sgRNAs inside cells and in the infectious progeny differentiate between an effect on viral genome replication, in which case the sgRNA abundance would change inside infected cells, and an effect after viral DNA replication, in which case the sgRNA abundance would not change significantly in the cells but will change in the infectious progeny. Indeed, the abundance of sgRNAs targeting RRM2 are reduced in cells, but do not change much further in the infectious progeny, illustrating RRM2 depletion mostly alters viral DNA replication (Figure 2f). In contrast, the sgRNAs targeting ARL6IP6 did not change drastically in cells but were significantly reduced in the infectious progeny showing ARL6IP6 depletion effects occur mostly post viral DNA replication (Figure 2i).Similarly, these distinct profiles were also observed for restriction factors (Supplementary figures 3a and 3b).

Interestingly, there was a small number of genes whose depletion showed opposing effects between the change in sgRNA abundance in cells as opposed to infectious progeny. The most striking example was NAA30, the catalytic subunit of the N-terminal acetyltransferase C (NatC) complex that catalyzes co-translational acetylation of methionine residues of specific proteins. Although NAA30 depletion led to reduction in DNA replication, there was a relative increase in the amount of infectious progeny (Figure 3a), suggesting NAA30 likely has pleiotropic effects on HCMV propagation and concurrently with its positive effects on DNA replication it has negative effects on later stages of the HCMV life cycle.

**Figure 3.**
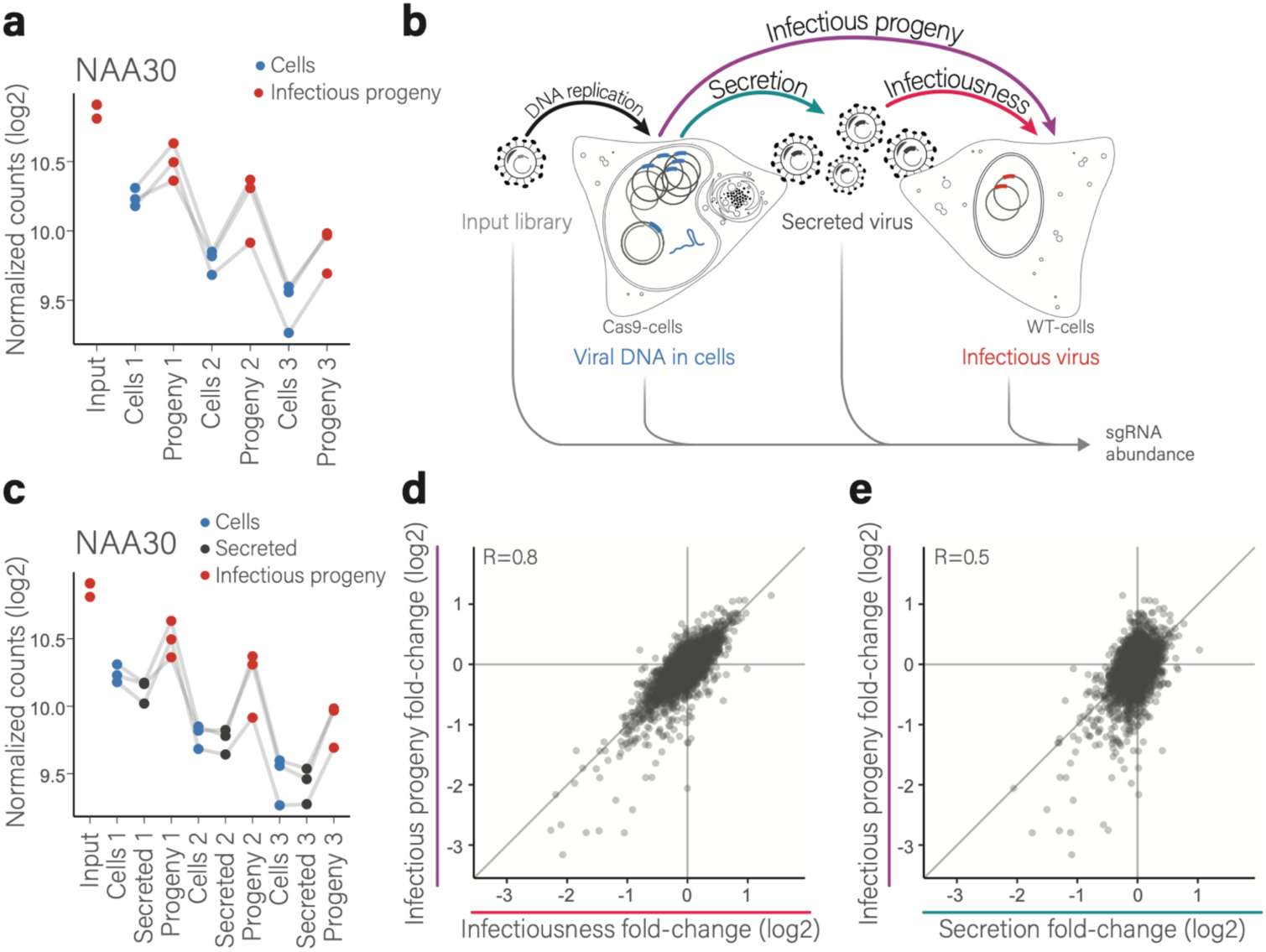
VECOS reports on the specific infection stage affected by perturbation. a. Relative levels of sgRNAs (geometric mean) targeting NAA30 at different stages along the screen (cells, blue; infectious progeny, red). The geometric means of sgRNAs targeting the gene are shown. Gray lines connect the values of the same biological replicate. b. Illustration of the different stages of HCMV infection as measured in the VECOS screen. DNA replication effects are reflected in changes between input virus and the measurements in cells (black arrow). Secretion effects are reflected as changes between the cells and the measurements of total secreted virus (cyan arrow). Infectiousness effects are reflected as the changes between total secreted virus to measurements of infectious virus (magenta arrow). Effects on total infectious progeny are the combined effect of secretion and infectiousness and are reflected in the change between sgRNA measurements in cells and infectious virus (purple arrow). c. Relative levels of sgRNAs (geometric means) targeting NAA30 along the screen (cells, blue; secreted particles, black; infectious progeny, red). Gray lines connect the values of the same biological replicate. d-e. Scatter plots showing the correlation between changes in secretion (d) or infectiousness (e) and changes in total infectious progeny at 10 dpi. Each point represents the change in one sgRNA in one replicate of one round of selection. Data is presented for significantly changing genes. Pearson’s R values are shown.

Changes in the load of infectious progeny post DNA replication could stem from effects on the physical amount of viral particles that are being secreted or may reflect changes in the quality or infectiousness of the secreted particles (Figure 3b). To quantitatively assess the contribution of these two processes, we additionally measured sgRNA abundance directly from the secreted viral particles (regardless of their infectivity potential), by pelletting viral particles from the supernatant and sequencing the sgRNAs (see Methods, supplementary figure 3c). For many of the hits which affected the amount of infectious progeny, sgRNA abundance in the secreted particles largely resembled the sgRNA abundance that was measured in cells (Figure 3c and Supplementary figure 3d-g). This suggests that for these genes particle secretion out of the cell is hardly affected by the perturbation and it is virion quality (infectiousness) that is the primary factor that affects the production of infectious progeny.

We next globally analyzed the contribution of changes in particle quantity, i.e. secreted particles (calculated from the difference in sgRNA abundance inside cells and in the secreted particles) and changes in particle quality, i.e. infectiousness (calculated from the difference in sgRNA abundance in secreted particles and in the infectious virus) to the total changes in infectious progeny (calculated from the difference in sgRNA abundance inside cells and in the infectious virus)(Figure 3b). Most of the changes in infectious progeny were explained by changes in particle quality whereas changes in particle quantity explain a smaller portion of the differences (R=0.8 and R=0.5, respectively, Figures 3d and 3e). Furthermore, for significant hits in the screen, the effect on infectiousness is on average 13-fold greater than the effect on secretion. Since we analyzed the sgRNAs in the cells, supernatant, and infectious particles, at 10 dpi, we seeked to verify that the apparent scarcity of perturbations that affect particle secretion is not due to nonspecific effect of cell death, at late time point of infection. We therefore repeated the VECOS screen in triplicates and collected the cells, supernatant and infectious particles at 5 dpi, when cells are still intact (Supplementary figure 3h). Similar to the 10 dpi screen, in the 5 dpi screen the vast majority of changes in the amount of infectious particles is explained by differences in the infectioness of the particles, and particle secretion has only a minor contribution (R=0.7 vs R=0.2, Supplementary figures 3i and 3j). Together, these analyses imply that most host genes that affect HCMV propagation post DNA replication actually influence particle quality and not the quantity of secreted viral particles.

These measurements of sgRNA distribution inside infected cells, in secreted particles, and in infectious progeny, provide a quantitative assessment on the effect of each gene on three stages in the viral life cycle: 1. Viral DNA replication (comparing sgRNAs in input viruses to sgRNAs in cells), 2. Secretion of viral particles (comparing sgRNA distribution in cells to sgRNAs distribution in secreted particles) and 3. Particle quality/ infectioness (by comparing sgRNA distribution in particles to sgRNAs distribution in infectious virus) (Figure 3b). Therefore, our data can be used to classify the effect of genes on these three steps in the infection cycle. We focused on 62 hits that showed statistically significant changes in one or more of these specific viral life cycle stages and we used hierarchical clustering to group these genes into 10 clusters according to the stages in which they affect infection. The resulting clusters included six clusters of dependency factors and four clusters of restriction factors. These clusters were further divided based on their effects on viral DNA replication, particle secretion and particle infectiousness (Figure 4a, supplementary figure 4a and supplementary table 3). Reassuringly, the two clusters that showed the strongest effects on viral DNA replication, which for HCMV occurs in the nucleus, were enriched with nuclear proteins (clusters 2 and 7, p-value = 0.046, hypergeometric test). On the other hand, clusters that include genes that have strong effects on infectiousness, were enriched in proteins with membrane association (clusters 4 and 8, p-value = 0.009, hypergeometric test), indicating particle infectiousness is often associated with membrane related functions (Figure 4a). Furthermore, profiles are consistent among individual sgRNA targeting the genes in each cluster (Supplementary figure 4b).

**Figure 4.**
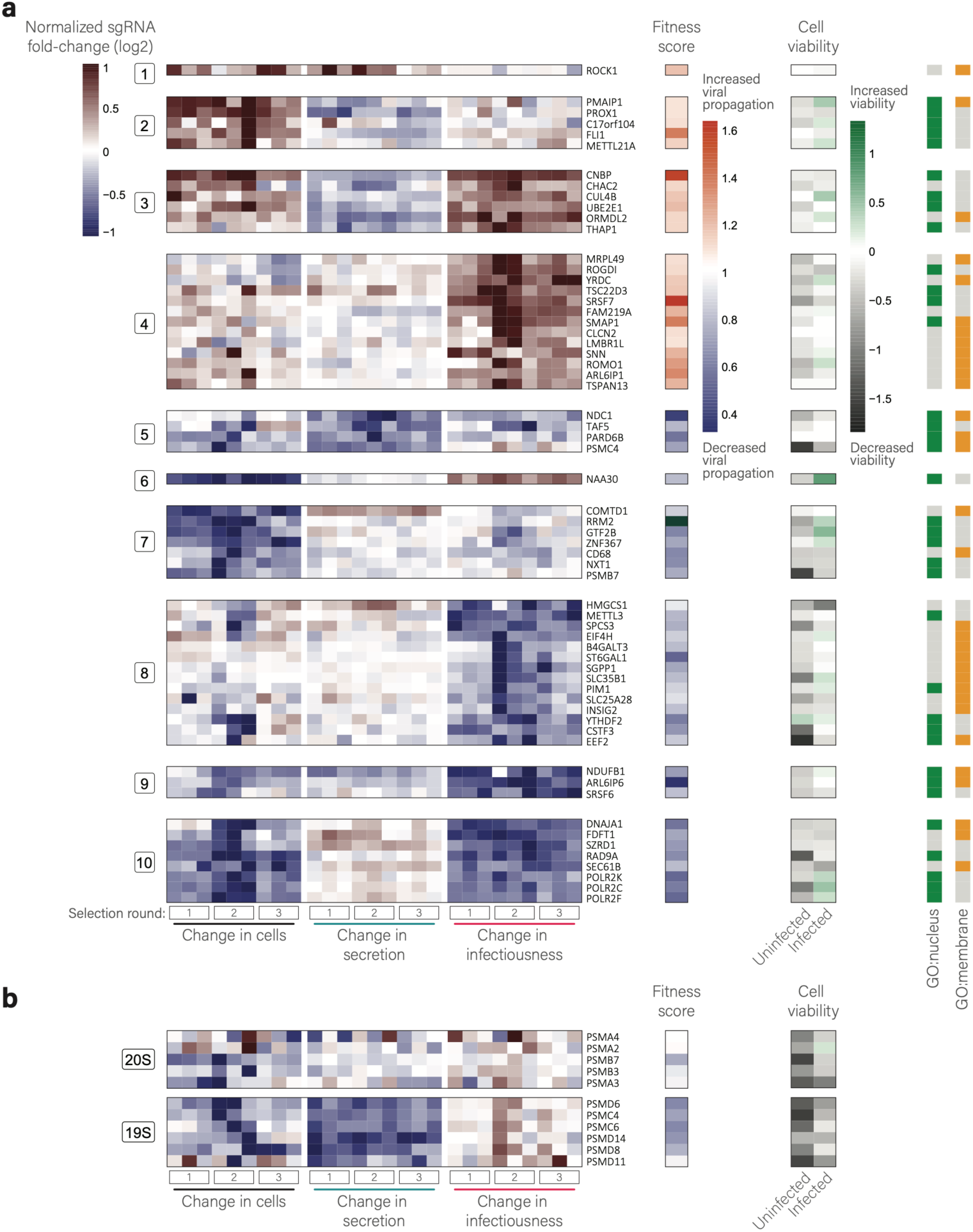
Dissection of host factors effects on three stages in HCMV life cycle. a. Heat map of fold-change in sgRNA abundance (normalized log2 transformed) of selected genes during different stages of viral infection. Colors represent the increase (red) or decrease (blue) in abundance of the sgRNAs targeting each gene. Fold-changes during viral replication in cells are shown on the left, fold-changes in secretion are shown in the middle and fold-changes in infectiousness are shown on the right, and for each gene. Values presented are of three biological replicates for each selection round (1,2, and 3). Annotations on the right show the fitness score, the cell viability effects of gene perturbations in uninfected and HCMV infected fibroblasts (Hein and Weissman 2021), and gene association with GO terms; nucleus (green) and membrane (yellow). b. Heat map of fold-change sgRNA abundance, fitness score and cell viability as described in a, for genes that encode for 26S proteasome subunits.

Interestingly, the two proteasome proteins that were included in this analysis were assigned to different clusters. Knockout of PSMB7, a component of the core 20S proteasome shows a pattern which is consistent with some reduction in viral DNA replication, maybe due to increased cell death. In contrast, the knockout of PSMC4, a component of the regulatory 19S subunit, mainly leads to a significant reduction in genome secretion (Figure 4a). To further explore these differences, we extended the analysis by plotting the profiles from the screen of all proteasome genes that were included and discovered a clear separation between the 20S and the 19S subunit components (Figure 4b). While knockout of both 20S and 19S (26S) components show a similar mild effect on fibroblast survival which is largely diminished when cells are infected, 19S subunits have a much stronger effect on HCMV propagation and only their depletion leads to a significant effect on the secretion of HCMV virions from the cell (Figure 4b). The 19S subunit of the proteasome has a regulatory activating effect on the function of the 20S core proteasome subunit, as well as additional potentially independent functions ^22,23^. Our results suggest the non-proteolytic function of the 19S proteasome is specifically important for HCMV propagation. More generally, this data illustrates that focused analysis on specific processes or complexes using VECOS can help to accurately define HCMV vulnerabilities to genetic or pharmacological interventions.

We next validated some of the phenotypes that came out of these clusters by generating KO cells by transducing cells with sgRNA and Cas9 prior to infection. Clusters 2 and 3 are composed of restriction factors that have an effect at the stage of viral genome replication. Indeed, compared to the control cells, knocking out FLI1, CNBP and CHAC2 prior to infection led to an increase in viral DNA replication and viral titers (Figure 5a and 5b). Depletion of genes from cluster 4 is predicted to specifically improve virion infectiousness. However, knocking out ROMO1, ARL6IP1 and SRSF7 prior to infection surprisingly led to reduced viral propagation (Supplementary figure 5a). In contrast, the phenotype observed in the screen was reproduced when using the VECOS system to express individual sgRNAs, where knock out occurs following infection. In this case virions budding out of Cas9 cells had significantly increased titers compared to virions budding out of control cells, and the effects on the amount of total secreted virus were much milder (Figure 5c-5f, and supplementary figure 5b and 5c). The difference between the results obtained from the different experimental systems suggest a time-dependent effect for these genes. Other genes predicted to modify virion infectiousness are the dependency factors grouped in cluster 8. Depletion of METTL3, HMGCS1, SGPP1 and ST6GAL1 prior to infection significantly reduced viral titers (Figure 5g and 5h), but had only minor effect on the amount of secreted virus (Figure 5i and 5j, and supplementary figure 5d and 5e), confirming that depletion of these four genes mostly reduce HCMV particle infectiousness.

**Figure 5.**
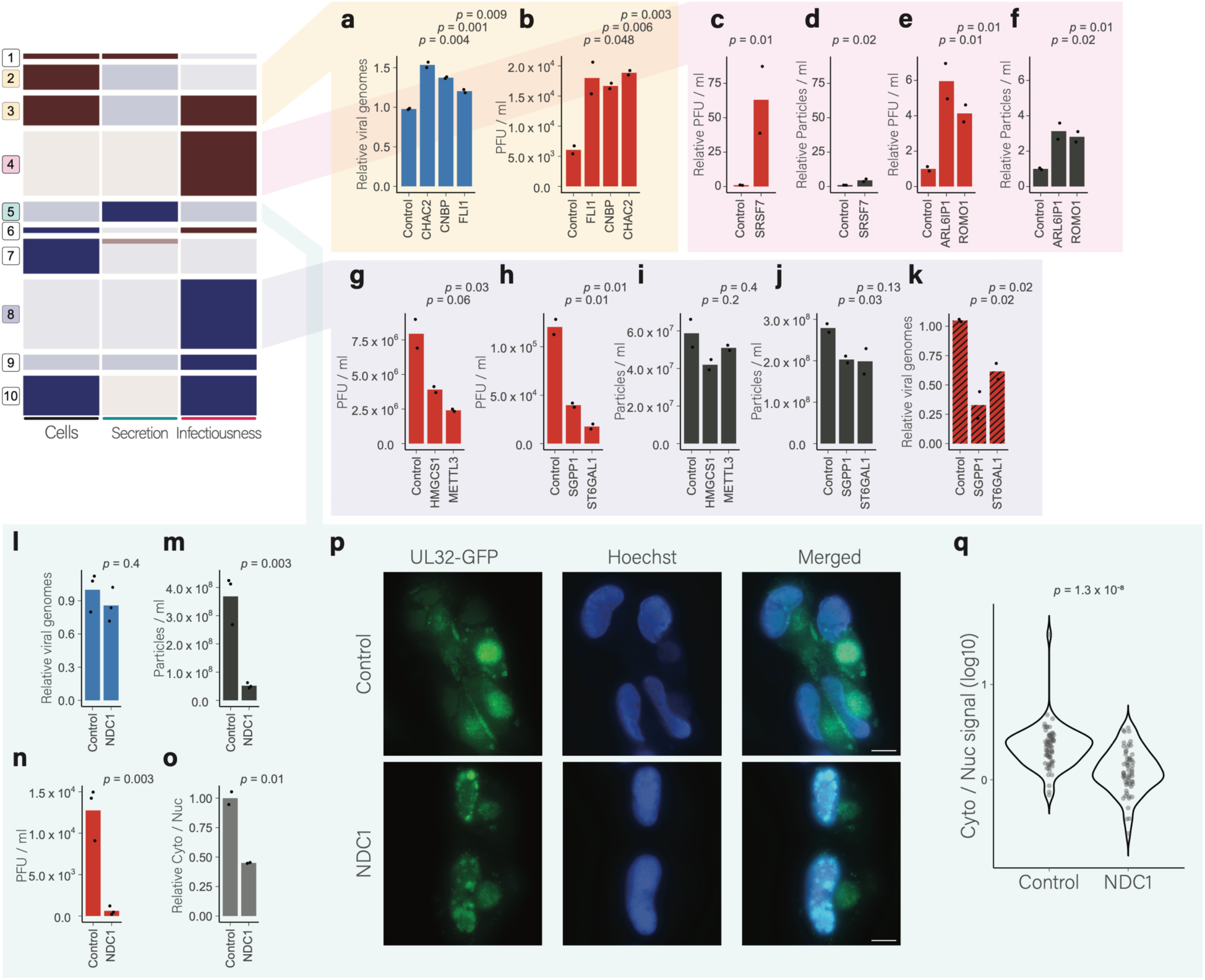
Validation of host factors effects on different stages of HCMV infection. a-n. Measurements of viral genome replication (a, l, blue bars), viral titers (b, c, d, g, h, n, red bars), secreted viral particles (e, f, i, j, m, dark gray bars), and virus entry (k, red striped bars) in various knockout fibroblasts (indicated in x-axis) compared to control. (a, l) DNA from knockout and control cells infected with HCMV was extracted at 3 dpi and the levels of host and viral DNA was measured by qPCR. Viral DNA levels were normalized to host DNA and to the control samples mean. (b, g, h, n) Viral supernatants were collected from knockout and control cells at 6 dpi and transferred to recipient wild-type fibroblasts. After 48 h, the recipient cells were analyzed by flow cytometry. PFU / ml were calculated from the percentages of GFP positive cells. (c,d,e,f) Viral supernatants were collected from Cas9- and control- (mCherry) expressing cells infected with sgRNA-encoding HCMV at 7 dpi. (c, e) Supernatants were transferred to recipient wild-type fibroblasts, and 48 h later, the recipient cells were analyzed by flow cytometry. Relative PFU / ml represents the ratio between supernatants from Cas9 and control cells. (d, f) Supernatants were boiled and SybrGold was used to stain DNA. The concentration of DNA containing particles was measured by small particle flow cytometry. Relative particles / ml represents the ratio between supernatants from Cas9 and control cells. (i, j, m) Viral supernatants were collected from knockout and control cells. Supernatants were boiled and SybrGold was used to stain DNA. The concentration of DNA containing particles was measured by small particle flow cytometry. (k) Viral supernatants collected from knockout and control cells were used to infect recipient wild-type fibroblasts. After 1 h incubation, extracellular virus was washed away, intracellular DNA was extracted from the cells at 5 hpi. The levels of host and viral DNA were measured by qPCR. o. Normalized ratios of viral DNA levels in cytosolic (Cyto) and nuclear (Nuc) fractions isolated from HCMV infected NDC1 knockout and control cells at 4 dpi (light gray bars). p. Fluorescence microscopy of control and NDC1 knockout fibroblasts infected with HCMV strain expressing GFP fused to UL32 (green) and nuclear Hoechst staining (blue) at 72 hpi. Scale bar is 10 μm. q. Distribution of cytosolic (Cyt) to nuclear (Nuc) ratios (log10) of GFP signal in control and NDC1 knockout cells. Points represent individual cells. In all plots, p-values were calculated using Student’s t-test. Bars represent the means of biological replicates.

ST6GAL1 Transfers sialic acid to galactose-containing acceptor substrates. SGPP1 catalyzes the degradation of Sphingosine-1-phosphate (S1P) via salvage and recycling of sphingosine into long-chain ceramides. We therefore hypothesized that reduction in particle infectivity in ST6GAL1 and SGPP1 KO cells may reflect changes in entry due to changes in viral envelope protein glycosylation or membrane lipid composition. To test whether effects on infectiousness come from differences in entry, we measured the entry of virions produced in SGPP1 and ST6GAL1 knockout cells. Intracellular DNA was extracted from infected cells at 5 hours post infection. Virions produced in SGPP1 or ST6GAL1 knockout cells entered cells less efficiently than virions produced in control cells (Figure 5k), indicating that these virions are at least partly defective in their ability to attach or fuse to cells.

Finally, we focused on NDC1, one of the few genes grouped in cluster 5, whose depletion led to reduction in viral particle secretion. NDC1 is a component of the nuclear pore complex (NPC) that plays a role in the assembly and insertion of NPC into the nuclear envelope. In agreement with the screen’s results, depletion of NDC1 had no drastic effect on viral DNA replication in cells (Figure 5l), but led to a significant reduction in viral DNA in the supernatant (Figure 5m and supplementary figure 5f) and to reduction in viral titers (Figure 5n), illustrating NDC1 indeed reduces the secretion of viral particles. Cytomegalovirus capsids are assembled in the nucleus and then cross the nuclear envelope in a unique process promoted by the viral nuclear egress complex. Given NDC1’s role in NPC assembly we tested whether NDC1 affects the translocation of viral capsids from the nucleus to the cytoplasm. Depletion of NDC1 resulted in lower relative abundance of viral genomes in the cytoplasmic fraction (Figure 5o) as well as nuclear retention of the capsid associated tegument protein, UL32 (Figure 5p and 5q), demonstrating disruption of viral capsid egress out of the nucleus. Thus, although the NPC is not thought to be involved in HCMV nuclear egress, NDC1, one of the NPC components, is used by the virus to facilitate this process.

Targeting of only one gene, ROCK1 (cluster 1) resulted in a significant increase in particle secretion. Interestingly, we previously showed that ROCK1 activity increases nuclear egress of HCMV particles to the cytoplasm ^11^. The observation that only few genes significantly affected viral particle secretion and from these the two high confidence hits that we analyzed actually affect HCMV egress out of the nucleus strongly imply that once encapsidated genomes make it to the cytosol, secretion out of the cells represents a default route and there is no single unique mechanism that regulates viral particle secretion.

In summary, we describe the establishment of VECOS, a new screening platform for dissecting viral-host interactions. VECOS provides superb sensitivity, due to direct probing of viral propagation instead of the cell’s state. This screening strategy also overcomes the bias towards perturbations that affect early events in infection, allowing discovery of critical novel factors in late stages of virus propagation. Moreover, VECOS facilitates in-depth classification of the stages in the virus life cycle that are affected by each gene perturbation. Applying this approach to HCMV infection resulted in a rich dataset of infection stage-specific host-virus dependencies and revealed that diverse genetic perturbations converge on affecting virion quality. These findings highlight that viral particle assembly and integrity, but not secretion, are critical stages in the HCMV cycle that are heavily reliant on cellular proteins.

Importantly, since VECOS relies on low multiplicity of infection, it could be extended to study factors important for HCMV propagation in various cell types, including cell models of latency and potentially more complex infection models such as organoids. More generally, our approach can be extended for studying additional herpesviruses and likely to other viruses ^24,25^, revolutionizing our ability to define viruses’ vulnerabilities to genetic or pharmacological interventions.

In sum, VECOS is a sensitive and versatile platform for high-resolution functional dissection of host-virus interactions that can serve as a transformative tool for designing more effective antiviral therapeutics.

## Materials and methods

### Cell lines and virus strains

Human foreskin fibroblasts (ATCC CRL-1634), W162 cells, and HEK293T cells were cultured in Dulbecco’s modified Eagle’s medium (DMEM, Biological Industries) supplemented with 10% (vol/vol) heat-inactivated fetal bovine serum (FBS, Life Technologies), 2 mM L-glutamine (Biological Industries), 0.1 mg/mL streptomycin and 100 U/mL penicillin (Biological Industries), at 37°C with 5% (vol/vol) CO2. HCMV strains used are Merlin UL32-GFP ^26^, and the viruses derived from the HCMV-GW (see below), as indicated for each experiment. The Merlin BAC encodes a complete wildtype HCMV genome with the exception of point mutations in RL13 and UL128, which are required for efficient propagation in fibroblasts ^27^. Adenovirus dl1014 mutant virus containing a deletion in the E4 locus ^28^ and the complementary W162 cell line ^29^ Iwere kindly provided by T. Kleinberger.

Fibroblasts were infected with HCMV by adding the virus to the cells and incubating with gentle shaking for one hour at room temperature before removing the virus containing supernatant and replacing it with fresh media. To propagate Adenovirus dl1014, W162 cells were infected with dl1014 at an MOI of 0.01 by incubating the cells with the virus with gentle shaking for 1.5 hours at room temperatures. The infected cells were collected 60 hpi and frozen at −80°C. Intracellular virus was released by adding 4-5 ml of serum free DMEM and performing three cycles of freeze-thaw by alternating the tubes between liquid nitrogen and a 37°C water bath. Cellular debris was then removed by centrifugation at 3,800 g for 20 minutes.

### Generation of lentiviruses for transduction

Lentiviruses were generated by co-transfection of vector constructs and second-generation packaging plasmids (psPAX2, Addgene no. 12260 and pMD2.G, Addgene no. 12259), using jetPEI DNA transfection reagent (Polyplus transfection), into HEK293T cells, according to the manufacturer’s instructions. At 48 h post transfection, supernatants were collected, purified, and filtered through a 0.45 μm polyvinylidene fluoride filter (Millex).

### Generation of plasmids

#### pDONR-sgRNA plasmid

The pDONR201 plasmid was modified for Gateway cloning of sgRNA constructs as follows. The U6-sgRNA construct was amplified from the pX330 plasmid (Addgene no. 42230) with addition of Gateway AttB1/2 recombination sites as flanking overhangs, using the primers GW-sgRNA-F and GW-sgRNA-R (Supplementary table 4).

The amplified product was combined with the pDONR201 plasmid in a gateway BP recombination reaction (Thermo-Fisher, catalog number 11789020) to get the U6-sgRNA cassette flanked by AttL1/2 sites in the pDONR201 plasmid. Next, the BpiI-FD (Thermo-Fisher) restriction sequence located on the pDONR201 plasmid outside of the inserted sgRNA construct was removed using the primers Bpi-del-F and Bpi-del-R (Supplementary table 4) in a quick change no kit cloning protocol (Source: Dylan Webster, Adapted from QuickChange II XL Site-Directed Mutagenesis Kit Protocol).

#### lentiCRISPRv2-dU6 plasmid

To introduce Cas9 into cells, the lentiCRISPRv2 plasmid ^30^ was modified by removing the U6-sgRNA cassette. The plasmid was digested with BsmBI (Thermo-Fisher, catalog number FD0454) and KpnI-HF (NEB) in Cutsmart buffer. Partially complementing oligos, U6-del-F and U6-del-R (Supplementary table 4) were annealed and phosphorylated using T4 PNK (NEB) and ligated into the restricted plasmid using T7 DNA ligase, as previously described for sgRNA cloning (Sanjana, Shalem, and Zhang 2014).

#### lentiCRISPRv2-2guide plasmid

To express two sgRNAs from one plasmid, the LentiCRISPRv2 plasmid was modified using restriction-free cloning to create a new plasmid called LentiCRISPRv2-2guide as follows. The cassette containing the sgRNA sites along with the U6 and H1 promoters as well as the sgRNA scaffolds was lifted from pDECKO_GFP plasmid ^31^ using primers, pDecko-GFP-RF-F and pDecko-GFP-RF-R (Supplementary table 4), which contain 5’ overhangs homologous to the LentiCRISPRv2 plasmid ^30^. The lifted fragment was then inserted into the LentiCRISPRv2 plasmid in place of the sgRNA cloning site. The PCR product was subsequently digested with DpnI and transformed into bacteria. For subsequent cloning of specific sgRNAs, the 2-sgRNA cassette was amplified from the LentiCRISPRv2-2guide plasmid using primers containing the desired sgRNA sequences (Guide2-cloning-F and Guide2-cloning-R, Supplementary table 4), and reinserted into the plasmid followed by DpnI digestion and transformation. For RRM2 knockout cells, the original lentiCRISPRv2 plasmid was used with one sgRNA cloned as previously described.

#### ARL6IP6-strepII over expression plasmid

ARL6IP6 was cloned into pLVX-Puro-TetONE-SARS-CoV-2-nsp1-2XStrep (kind gift from N. Krogan, UCSF) in place of the SARS-CoV-2-nsp1 cassette. ARL6IP6 coding sequence was amplified from cDNA with primers containing flanking regions homologous to the vector (Supplementary Table 4) and the plasmids were amplified with the appropriate primers (Supplementary Table 4). The amplified PCR fragments were cleaned using a gel extraction kit (Promega) according to the manufacturer’s protocol and used for cloning using a Gibson assembly protocol. The expression of ARL6IP6 was induced by adding doxycycline 24 hours prior to infection and removing it at the time of infection.

### HCMV-GW construction

Gateway recombination sites flanking a ccdB expression cassette were engineered into the Merlin BAC in place of the gene RL13, which must be mutated to permit efficient growth *in vitro*. The parental BAC also contained a point mutation in UL128 to enable efficient dissemination, and a P2A-GFP cassette linked to the C-terminus of IE2 to track infection. Firstly, a cassette encoding rpsL, lacZa, and kanamycin resistance, was amplified using primers Kan_ins_RL13_F and Kan_ins_RL13_R (Supplementary table 4). Then inserted into the RL13 ORF by recombineering, as previously described ^32^.This BAC was then transferred into gBRed gyrA462 bacteria, which contain a mutation rendering them resistant to the lethal effects of the ccdB gene, and were a gift from Francis Stewart (Technische Universitat Dresden ^33^). Once transferred, a cassette encoding AttR, cmR, ccdB, and AttR sites was amplified from pLenti6-V5-Dest-Empty using primers GW_ins_RL13_F and GW_ins_RL13_R (Supplementary table 4). This PCR product was used to replace the rpsL/LacZa/KmR cassette by recombineering, before plating on media containing Streptomycin as counterselection against the original cassette. All constructs were verified by Sanger sequencing across the modified site.

### Preparation of inactivated adenovirus particles

Adenovirus particles for Adenofection were prepared as previously described ^6,7^, with a few small adjustments. Adenovirus particles were purified using isopycnic cesium chloride (CsCl) gradient centrifugation, protocol adjusted from JoVE ^34^. CsCl solutions were prepared in three different concentrations. To achieve density of 1.5 g/cm^3^, 45.4 g of CsCl were dissolved in 54.6 ml of water, for 1.35 g/cm^3^, 35.2 g of CsCl were dissolved in 64.8 ml of water, and for 1.25 g/cm^3^, 27 g of CsCl were dissolved in 73 ml of water.

For the first gradient, CsCl step gradients were prepared in one clear ultracentrifuge tube (Beckman-Coulter, catalog number 344059) by slowly pipetting the solutions in the following order: 0.5 ml of 1.5 g/cm^3^ CsCl solution, 3 ml of 1.35 g/cm^3^ CsCl solution, and 3.5 ml of 1.25 g/cm^3^ CsCl solution. The gradient was overlaid with the virus supernatant, and centrifuged in an ultracentrifuge using a swing out rotor (SW-41 Ti) at 12°C for at least 2 hours at 226,000 g with slow acceleration and deceleration. The virus-containing band (lower bands) was identified by light scattering and collected by puncturing the tube with a hollow needle (about one ml collected volume). For the second gradient, the collected virus was transferred with a clean pipette tip into a sterile 50 ml tube, and 12 ml of 1.35 g/cm^3^ CsCl solution was added, and mixed carefully. The mixture was transferred to ultracentrifuge tubes, and centrifuged overnight at 12°C at 226,000 g with slow acceleration and deceleration. The virus-containing band (lower bands) was identified by light scattering and collected by puncturing the tube with a hollow needle (about 1 ml collected volume). The samples were desalted using PD-10 columns (GE healthcare, catalog number 17085101), 3.5ml elution buffer (HBS/40% glycerol, HBS is 20 mM HEPES, 150 mM NaCl pH 7.4) was added and fractions of 0.5 ml each were collected into separate tubes. Virus-containing fractions were identified by measuring protein concentrations with the Pierce BCA protein assay kit (Thermo-Fisher), and protein-positive fractions were pooled. To inactivate the virus, 8-methoxy-psoralen (Sigma, M-3501) was added to a final concentration of 0.33 mg/mL, the sample was transferred on ice and exposed to UV light (366 nm) for 30 min, with rotation of the dish every 10 minutes. The inactivated virus was desalted again as described above, and the concentration of Adenovirus particles in the desalted samples was determined by calculating 0.3 mg/mL of (capsid) protein being equivalent to 1 × 10^12^ particles/ml.

### Design of host targeting library

Genes targeted in the screen were selected based on mRNA expression data during HCMV infection ^5^, by choosing the 2,000 genes with the highest fold-change between the uninfected and the 72 hpi samples. Additional 240 genes of interest were added manually (Interferon stimulated genes and other genes of interest). For each gene, four sgRNAs from the published Brunello library ^35^ were included in the library.

### Cloning of VEKOS libraries and individual sgRNA viruses

VEKOS libraries and individual sgRNA viruses were constructed in three steps, while maintaining 10X coverage of the library at each step of its construction.

#### Step 1 – pDONR cloning

The sgRNA library with flanking homology regions (see below) was synthesized by TWIST Bioscience, amplified, purified, and cloned using Gibson assembly as a pool into a the pDONR-sgRNA plasmid (a modified pDONR201 that contains the U6 promoter and sgRNA construct between the attL1/2 recombination sites as described above). Individual sgRNAs were cloned into pDONR-sgRNA with a restriction-ligation protocol. Oligos with flanking overhangs were synthesized by Sigma-Aldrich, and cloned following annealing and phosphorylation.

Oligo design with flanking homology regions for library cloning, N represent the gRNA sequence:

5’ - GGAAAGGACGAAACACCG - NNNNNNNNNNNNNNNNNNNN –

TTTTAGAGCTAGAAATAGCAAGTTAAAATAAGGC – 3’

Oligo design with overhangs for single sgRNA cloning, N represent the gRNA sequence and its reverse complement:

5’ – CACCGNNNNNNNNNNNNNNNNNNN – 3’

3’ – CNNNNNNNNNNNNNNNNNNNCAAA – 5’

#### Step 2 – BAC cloning

The library was cloned into an HCMV BAC containing a cassette of the lethal CcdB gene flanked by attR1/2 recombination sites, located in place of the RL13 gene that was deleted (see above). The Donor library or individual sgRNA plasmids were linearized using the NheI-HF enzyme (NEB) in a 50 µl reaction using 6 µg plasmid DNA and 1 µl enzyme, and the reaction was purified using Wizard SV Gel and PCR Clean-Up System (Promega). The linearized Donor was combined with the HCMV GW BAC in a Gateway reaction with LR Clonase II Plus enzyme (Thermo-Fisher, catalog number 12538120) according to the manufacturer’s instructions, with 7.3 µl BAC DNA suspended in TE and 0.7ul linearized plasmid library and an overnight incubation at 25°C. The reaction was transformed into electro-competent bacteria prepared using an improved protocol as described in Nováková et al. ^36^ plated on warm Chloramphenicol-LB-agar plates, and grown at 30°C for two days (small colonies were observed about 24 hours after plating). The cloned BAC DNA was extracted using NucleoBond Xtra BAC kit for large construct plasmid DNA (Macherey-Nagel, catalog number 740436.25). For library cloning, all the bacterial colonies were collected by washing the agar plates with 6 ml of LB or PBS, scraping the colonies, and collecting the liquid into a 50 ml falcon tubes. Each plate was washed twice, and the bacteria were centrifuged at > 1,500 g for > 10 minutes to collect into a pellet. To extract BAC from the bacterial colonies, 6 NucleoBond columns were used.

#### Step 3 – Viral stock reconstitution

The BAC library was introduced into fibroblasts using the Adenofection method ^7^ . For Adenofection of one six-well of fibroblasts (∼300,000 cells), 1 µg of BAC DNA was carefully diluted in 250 µL of sterile HBS (20 mM HEPES, 150 mM NaCl pH 7.4), and 15 µL of 10 mM PEI 2000 stock solution (polyethylenimine MW 2000, Sigma, Cat. No. 40870-0, 0.9 mg/ml H2O), mixed before usage, was diluted in 250 µL of HBS and mixed vigorously again. The diluted PEI 2000 solution was slowly added to the diluted DNA, with constant gentle flicking of the tube, and the sample was incubated for 20 min at room temperature. The mixture was added with 3 × 10^9^ particles of inactivated adenovirus, and the sample was incubated at room temperature for an additional 20 min. Fibroblasts cultured in six-well plates were washed with PBS, and 1.5 ml of serum-free DMEM containing polymyxin B (30 µg/mL, Sigma-Aldrich, catalog number P4932) was added to each well. The transfection complexes were added to the cells by dripping and the cells were allowed to incubate for 5 hours at 37°C 5% (vol/vol) CO2. Next, the cells were washed twice with PBS and further cultivated in complete growth medium with polymyxin B (30 µg/mL). One week later, cells were harvested with Trypsin and re-seeded on top of fresh fibroblasts (50-70% confluent) in T175 flasks at a ratio of two Adenofected 6-wells to one flask, and the supernatant from the Adenofected wells was added to the flask as well. Following 5-6 days the media was changed, and after additional 8-9 days the media with the reconstituted virus was collected, aliquoted and frozen at −80°C.

### VECOS screens

Cells were transduced with lentiCRISPRv2-dU6 lentivector to express Cas9 (modified lentiCRISPRv2 with the U6-sgRNA sequence deleted as described above) that was titrated for the minimal concentration that allows most cells to be transduced (∼MOI=3). Positive cells were selected using puromycin (1.75 μg/ml) for 2-3 days, and the Cas9 cells were infected with the VEKOS library, at a low multiplicity of infection (MOI 0.05). The cells and media containing progeny virus were collected 10 days post infection and frozen at −20°C and −80°C, respectively. For multiple passages, progeny viruses were titrated and used to infect fresh Cas9 cells, repeating the process described above. To sequence the sgRNA distribution in the input library in infectious progeny of each selection round, fresh fibroblasts, not expressing Cas9, were infected with a high MOI with the original VEKOS library virus or virus supernatant collected from the screen, and the cells were collected 2-3 dpi and frozen at −20°C. To sequence the sgRNA distribution in secreted viruses, the remaining virus containing media was collected and centrifuged for 20 minutes at 6,000 g to remove cell debris. The supernatant was transferred to new bottles and centrifuged for 3.5 hours at 13,000g.

DNA from cells and virus pellets was extracted using a published protocol ^37^. The viral and cell pellets of each replicate were resuspended in 300 µl and 6 ml of NK buffer, respectively, and DNA extraction was continued normally with volumes adjusted to the 300 µl start, and GlycoBlue™ (Thermo-Fisher, catalog number AM9516) was added at the stage of DNA precipitation.

Finally, sgRNAs were PCR amplified with primers containing Illumina sequences as overhangs (Supplementary table 4). The reactions were performed at a 50 µl volume with 0.5 nM of each primer (forward and reverse), 25 µl NEBNext High-Fidelity 2X PCR Master Mix (NEB, catalog number M0541S). In each 50 µl reaction, 2.5 µg DNA was used as template in amplification of DNA extracted from cells, and 500 ng DNA amplification of DNA extracted from secreted virions. PCR reaction conditions included denaturation at 98°C for 2 minutes, and repeating cycles of annealing and elongation at 98°C for 10 seconds, 65°C for 10 seconds and 72°C for 30 seconds. These were repeated for 18 cycles for DNA extracted from cells and 20 cycles for DNA extracted from secreted virions. At the end of the cycles, final extension was performed at 72°C for 5 minutes.

The PCR products were purified using a two-sided SPRI beads cleanup at bead ratios of 0.5X and 0.9X (AMPure XP, Beckman Coulter), and their concentration was measured using high sensitivity D1000 DNA tapestation (Agilent).

### Quantification of sgRNAs and Identification of hits from VECOS screen

MAGeCK v0.5.6 ^10^ was used to count sgRNA from FASTQ files and for initial analysis of selection effects using the robust rank aggregation (RRA) algorithm with normalization to total reads. Directed FDR and RRA scores for genes were calculated by taking the enrichment MAGeCK FDR and RRA scores for genes with a positive fold-change from control and the depletion MAGeCK FDR and RRA score for genes with a negative or zero-fold change.

To analyze and incorporate the selection effects on sgRNAs and genes along the entire screen and selection stages (cells and progeny) into one score, a mixed model regression analysis was performed as follows. First, sgRNAs with counts lower than 20 in any of the input samples were filtered out. Next, sgRNA counts were normalized to the median count of control non-targeting sgRNAs. This normalization is based on the biological expectation that control sgRNAs are not under selection pressure and that changes in their abundance are due to random drift. By normalizing to the median of control sgRNAs we also overcome potential composition bias that may be caused with normalization to total. Finally, the normalized counts were converted to pseudo-counts with an offset of +1, and log2 transformed. For each gene, A linear mixed-model regression model was used ^38^. The normalized sgRNA abundance was used as the dependent variable, the selection round number and selection stage (cells and progeny) as the fixed effect variables, and biological replicate number and sgRNA identity as random effect variables, as described in the formula below.

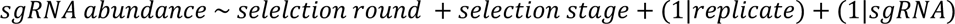

Including replicate number and sgRNA as random effects allows the regression model to treat the values from each sgRNA in each replicate as paired, and combine all values for a gene into one result. Selection round number was defined as −3 for the input samples, −2, 0, 2 for the cells samples in rounds 1, 2 and 3, respectively, and −1, 1 and 3 for the infectious progeny samples in rounds 1, 2, and 3, respectively. Since the regression model includes the selection stage, it produces (per gene) one slope value, and an intercept offset that reflects the differences between values at the cell stage and at the infectious progeny stage. The slope corresponds to the log2 fold-change from each selection step to the next. We defined the gene fitness score as 2^!∗#$%&’^, which corresponds to the effect size of a gene perturbation during one complete selection round (from input to infectious progeny). The regression test also provided a significance p-value, which was adjusted to multiple comparisons using the FDR method.

Thresholds for defining significantly changing genes were set by running the regression on 1000 simulated non-targeting genes. Each simulated gene was defined by selecting 4 random non-targeting sgRNAs. Significance and effect size thresholds were set as the values that allow 5% of the non-targeting genes to pass each threshold, separately. Next, the slope was tested for each sgRNA individually using the same regression approach with the formula listed below.

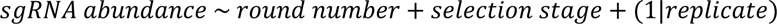

Genes were filtered out if they had less than two sgRNAs that had a significant (FDR<0.05) slope in the direction that matches that of the gene.

To identify genes with significant effect in individual stages of viral replication we repeated the regression with the following modifications, as seen in the formula below. The round number was set as the categorical random effect variable (pairing all samples of each round), and the two compared selection stages were set as the categorical fixed effect variables. This was done on all 3 pairs of selection stages, cells, secreted, and infectious virions.

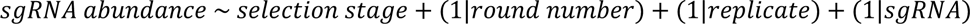

Genes were chosen as significantly changing in a specific selection stage if the FDR of the slope was below 0.1 in at least one of the stages. Genes that were not identified as hits in the main regression analysis were filtered out.

### Gene clustering based on different viral propagation stage effects

To analyze perturbation effects on different stages in viral propagation we focused on the genes that had significant effects on individual selection stages (replication, secretion or infectiousness), and the geometric means of sgRNA abundance for each gene were calculated. Next, the log2 difference between the values in each selection stage and the previous stage was calculated (e.g. log2(secreted - cells) for gene X of round 1 replicate A), resulting in scores for changes in cells, in secretion and in infectiousness, for each replicate in each selection round (in total 27 values for each gene). Next, the values were normalized across samples by dividing by the absolute maximum, so that the strongest change in each gene was normalized to either 1 or −1, depending on the direction of the effect. For clustering, the normalized values were averaged across biological replicates and selection rounds resulting in three values for each gene (cells, secretion and infectiousness). The genes were clustered based on the averaged normalized values using hierarchical clustering with the ward D2 algorithm, and visualized using the pheatmap package v1.0.12^39^. Genes were annotated as nucleus or membrane associated based on the Gene Ontology cellular component terms GO:0005634 and GO:0016020, respectively ^40,41^.

### Genomic DNA sequencing and analysis

To analyze the accumulation of mutations in genes targeted with VEKOS, Cas9 expressing fibroblasts were infected with VEKOS HCMV encoding for one sgRNA at about MOI 1, targeting either RRM2 or TRIP10 (Supplementary table 5). The cells were collected at different time points post infection and DNA was extracted (see above). The targeted genomic locus targeted in each sample was PCR amplified using the primers flanking the cut region (RRM2-F, RRM2-R, TRIP10-F, TRIP10-R, Supplementary table 4) that include overhangs complementary to Illumina sequences, and the product was purified with SPRI beads. Full Illumina adaptor sequences and barcodes were added in a second PCR reaction.

The library was sequenced with 150 cycles from each side and the sequences were aligned to the human genome using STAR v2.5.3a ^42^ with the following flags: scoreDelBase 0, scoreDelOpen 0, scoreInsOpen 0, scoreInsBase 0, alignIntronMin 300, alignIntronMax 299. The indel rate was calculated as the fraction of reads containing insertions or deletions based on the CIGAR sequence.

### Immunoblot analysis

Cells were lysed using ice-cold RIPA buffer (150 mM NaCl, 1% Triton X-100, 0.5% Na deoxycholate, 50 mM Tris-HCl [pH 8] and 0.1% (w/v) SDS) supplemented with a protease inhibitor cocktail (Sigma-Aldrich) and frozen at −80°C. Lysates were later thawed on ice and cleared by centrifugation at 4°C for 10 minutes at 20,800 x g. Proteins were separated on Bolt 4–12% Bis-Tris Plus polyacrylamide gels (Thermo-Fisher) and blotted onto nitrocellulose membranes. The membranes were blocked with Odyssey Blocking Buffer (Li-COR) mixed 1:1 with PBS, and immunoblotted with primary antibodies (antibody dilution indicated below, in TBST, 5% BSA and 0.05% (w/v) NaN3) for overnight at 4°C (TBST, 150mM NaCl, 50 mM Tris-HCl pH7.5 and 0.1% (v/v) Tween). This was followed by three washes with TBST. The membranes were probed with secondary antibodies for 1 hour at room temperature and washed three times with TBST. Fluorescent signal was acquired using an Odyssey CLx (LI-COR). The primary antibodies used were rabbit anti GAPDH (catalog number 2118S, cell signaling, 1:1,000), rabbit anti RRM2 (catalog number 65939, cell signaling, 1:1,000), mouse anti TRIP10 (catalog number 612556, BD bioscience, 1:500), and UL44 (Virusys, CA006). The secondary antibodies (1:20,000 in TBST and 5% (w/v) skimmed milk powder) used were as follows: IRDye 800CW goat anti-rabbit (Li-COR LIC-926-32211), IRDye 800CW goat anti-mouse (Li-COR, LIC-926-32210).

### Validation of screen results in knockout cells

To create individual gene knockout cells, fibroblasts were transduced with lentivirus expressing two sgRNA and Cas9 (lentiCRISPRv2-2guide) and were puromycin-selected (1.75 µg/ml) for 2 days. After selection, cells were split at a ratio of 1:2 and allowed to grow to full confluency before HCMV infection that was followed by different downstream analysis. In each experiment, three genes that showed no effect on infection in the screen were used as controls (CARD10, TRIP10 and IGSF8) exhibited similar behavior, and one is shown. All sgRNA used in individual assays are detailed in supplementary table 5.

### Immunofluorescence

Cells were plated on Ibidi slides, fixed in 4% paraformaldehyde for 10 min, permeabilized with 0.25% Tween20 and 10% normal goat serum (NGS) in PBS for 10 min, and then blocked with 0.2% Tween20 and 10% NGS in PBS for 1 hour. Detection of the strep-tagged ARL6IP6 was done with Strep-TactinXT DY-649 (IBA-lifesciences), detection of calnexin was done using rabbit calnexin polyclonal antibody (catalog number 10427-2-AP, proteintech) followed by donkey anti-rabbit–Rhodamine Red-X (Jackson Immunoresearch). Nuclei detection was done with Hoechst 33258. Imaging was performed on an AxioObserver Z1 wide-field microscope using a ×63 oil objective and Axiocam 506 mono camera using ZEN software.

### Viral particle flow cytometry

Viral supernatants were diluted 1:100 in PBS, and 5 µl were added to a 195 µl SybrGold solution (Invitrogen, catalog number S11494, diluted 1:10,000 in 0.2 µm filtered Tris-EDTA), following a 20 minute incubation at 80°C, while avoiding light exposure, to stain DNA. Samples were then analyzed by flow cytometry using a CytoFLEX S machine with a small particle detector (Beckman Coulter), and the concentration of the SybrGold positive population defined as virus particles (VP) was measured.

### qPCR of host and viral genomes

To extract DNA for quantitative real-time PCR (qPCR), cells were washed in PBS and lysed in a buffer mix containing a 1:1 ratio of buffer A (100mM KCl, 10mM Tris-HCl pH 8.3, 2.5mM MgCl) and buffer B (10mMTris-HCl pH 8.3, 2.5mM MgCl, 1% Tween20, 1% NP-40), with addition of 0.4 mg/ml proteinase K (Invitrogen, catalog number 25530049). The lysate was incubated for 1 hour at 60°C and then 10 minutes at 95°C. To extract DNA from viral supernatants, 0.4 mg/ml proteinase K was added and the supernatant was incubated for 1 hour at 60°C and then 10 minutes at 95°C.

Quantitative real-time PCR was performed using the SYBR Green PCR master-mix (ABI) on a QuantStudio 12K Flex Real-Time PCR System (Life Technologies) with primers targeting the human genome or the viral genome (human B2M-F, B2M-R, viral UL44-F, UL44-R, Supplementary table 4).

### Transfer assay (progeny)

Viral supernatants were collected from cells 7 dpi, and kept at −80°C or used immediately. Fresh fibroblasts were infected with the collected supernatants, and analyzed by flow cytometry 2-3 dpi to quantify GFP positive cells (infected cells). MOI was calculated as −ln(1-n), where n is the proportion of infected cells. Plaque forming units (PFU) per ml was calculated by multiplying the MOI by the number of cells in the infected well and dividing by infection volume in ml.

### Viral entry assay

Viral supernatants collected from knockout cells at 7 dpi were used to infect fresh fibroblasts. Following one hour of incubation with the virus, the cells were washed once with a citric acid buffer ^43,44^ (pH3, 40 mM citric acid, 10 mM KCl, 135 mM NaCl) for 50 seconds, and then briefly twice in PBS. Cells were lysed 5 hpi and host and viral DNA were quantified using qPCR.

### UL32GFP imaging and quantification

Cells were infected with Merlin UL32-GFP on Ibidi slides. At 4 dpi cells were fixed in 4% paraformaldehyde for 10 min, permeabilized with 0.1% triton X-100 and stained with Hoechst 33258. Imaging was performed on an AxioObserver Z1 wide-field microscope using a ×63 oil objective and Axiocam 506 mono camera using ZEN software. Quantification of cyto/nuc signal was calculated for each cell on ImageJ by defining the cell area using the brightfield channel and the area of the nucleus by using the Hoechst channel and quantifying the level of GFP (fused to UL32) in the nucleus area and in the cell area minus the nucleus area.

### Cellular Fractionation for quantification of viral genomes

Infected Cells were washed in cold PBS, and resuspended in 150µl buffer A (15 mM Tris-HCl pH 8, 15 mM NaCl, 60 mM KCL, 1 mM EDTA, 0.5 mM EGTA, 0.5 mM Spermidine), supplemented with 150 µl 2X lysis buffer (15 mM Tris-HCl pH 8, 15 mM NaCl, 60mM KCL, 1 mM EDTA, 0.5 mM EGTA, 0.5 mM Spermidine, 0.5% NP-40), mixed gently and incubated for 2 minutes on ice. The extract was centrifuged for 5 min at 400 g in a cold centrifuge, the supernatant was transferred to a new tube and centrifuged again for 1 min at 500 g in a cold centrifuge. 200µl of the supernatant (cytoplasmic fraction) was transferred to a new tube and 0.4mg/ml proteinase K was added. Residual supernatant was removed from the nuclear pellet and 1ml RLN (50 mM Tris-HCl pH 8, 140 mM NaCl, 1.5 mM MgCl2, 10 mM EDTA, 0.5% NP-40) was added. Following 5 minute incubation on ice, cells were centrifuged for 5 min at 500 g in a cold centrifuge and the supernatant was removed. 200µl of DNA lysis buffer (10 mM Tris-HCl pH 8, 50 mM KCl, 2.5 mM MgCl, 0.5% Tween20, 0.5% NP-40, 0.4mg/ml Proteinase K) was added to the nuclear pellet. All samples were incubated at 60°C for 1 hour and boiled at 95°C for 10 minutes prior to qPCR.

### General statistics

Statistical significance of individual assay results was calculated by performing two-sided Student’s t-tests assuming equal variance, or by performing hypergeometric tests (as indicated for each experiment). All plots and statistical tests were done using R 4.2.0 ^45,46^

## Acknowledgements

We thank Igor Ulitsky, Schraga Schwartz and the members of the Stern-Ginossar lab, for critical reading of the manuscript. We thank Hadas Hezroni-Bravyi for guidance during early protocol establishment and the Weizmann flow cytometry unit for technical assistance. This study was supported by MRC (MR/S00971X/1) and Wellcome Trust (226615/Z/22/Z) and European Research Council consolidator grant to N.S-G (CoG-2019-864012).

