## Supplementary figures and images for "A virally encoded high resolution screen of cytomegalovirus host dependencies"

### Supplementary figure 1

Supplementary figure 1

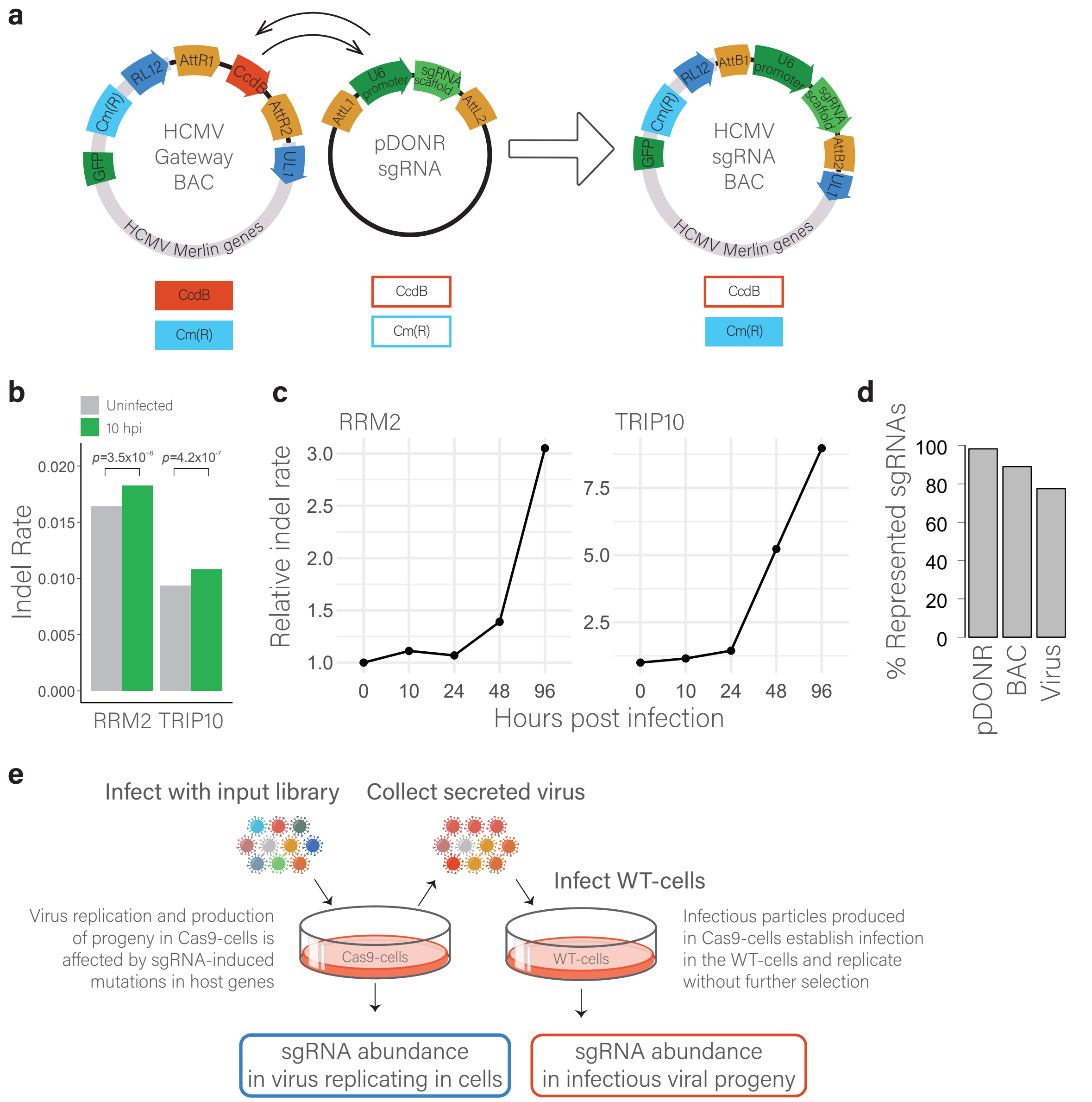

### Supplementary figure 2

Supplementary figure 2

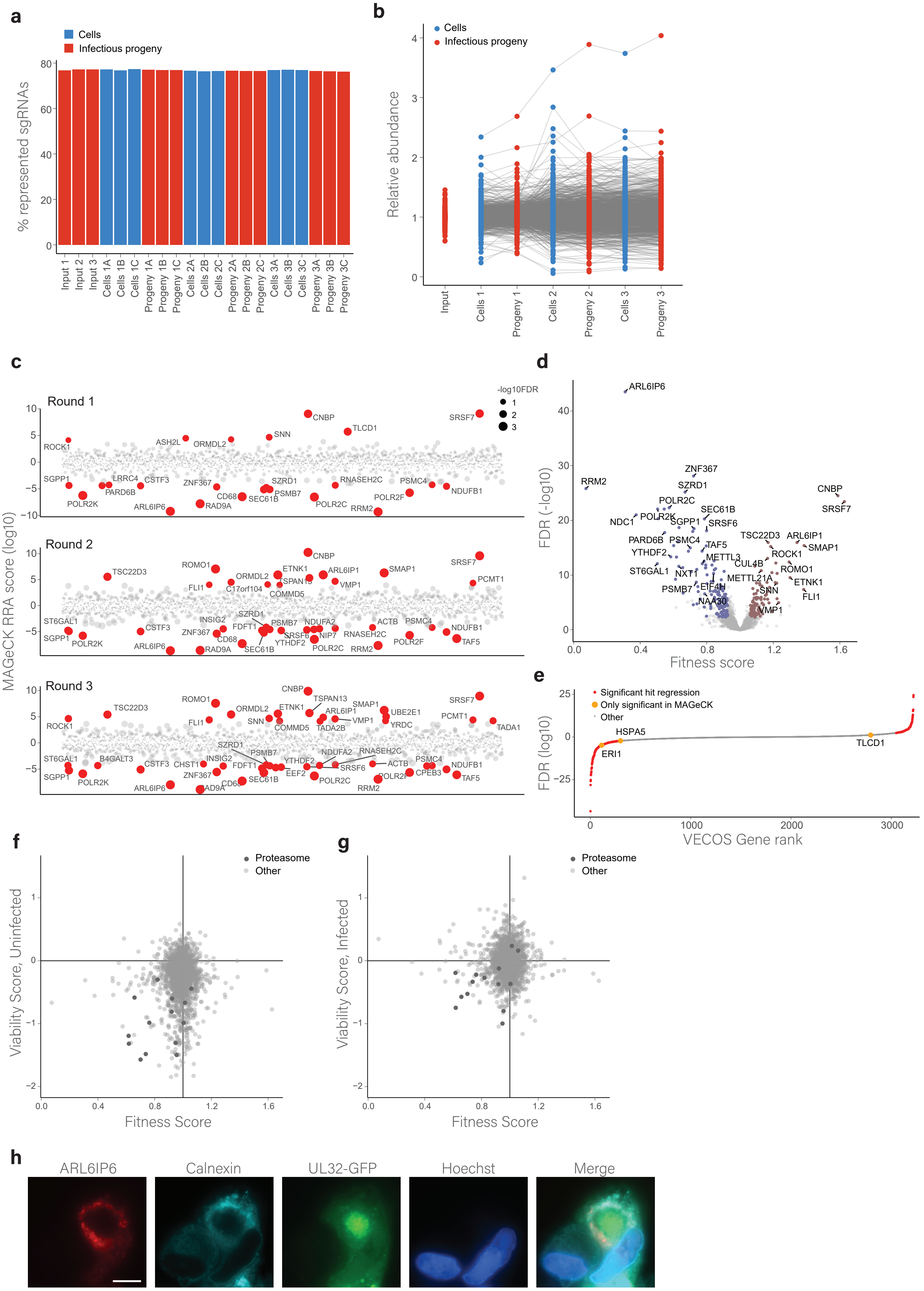

### Supplementary figure 3

#### Supplementary figure 3

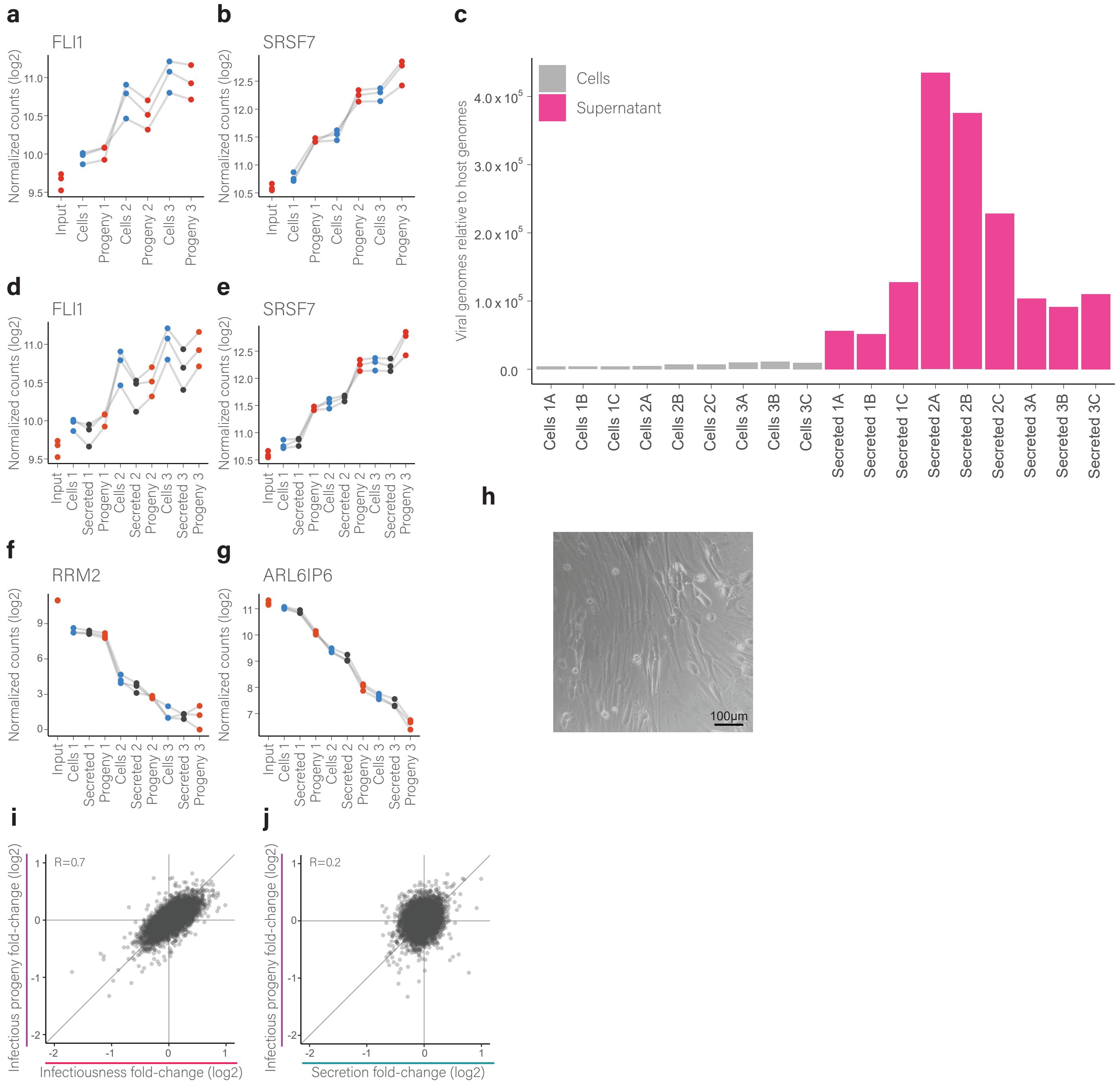

### Supplementary figure 4

Supplementary figure 4

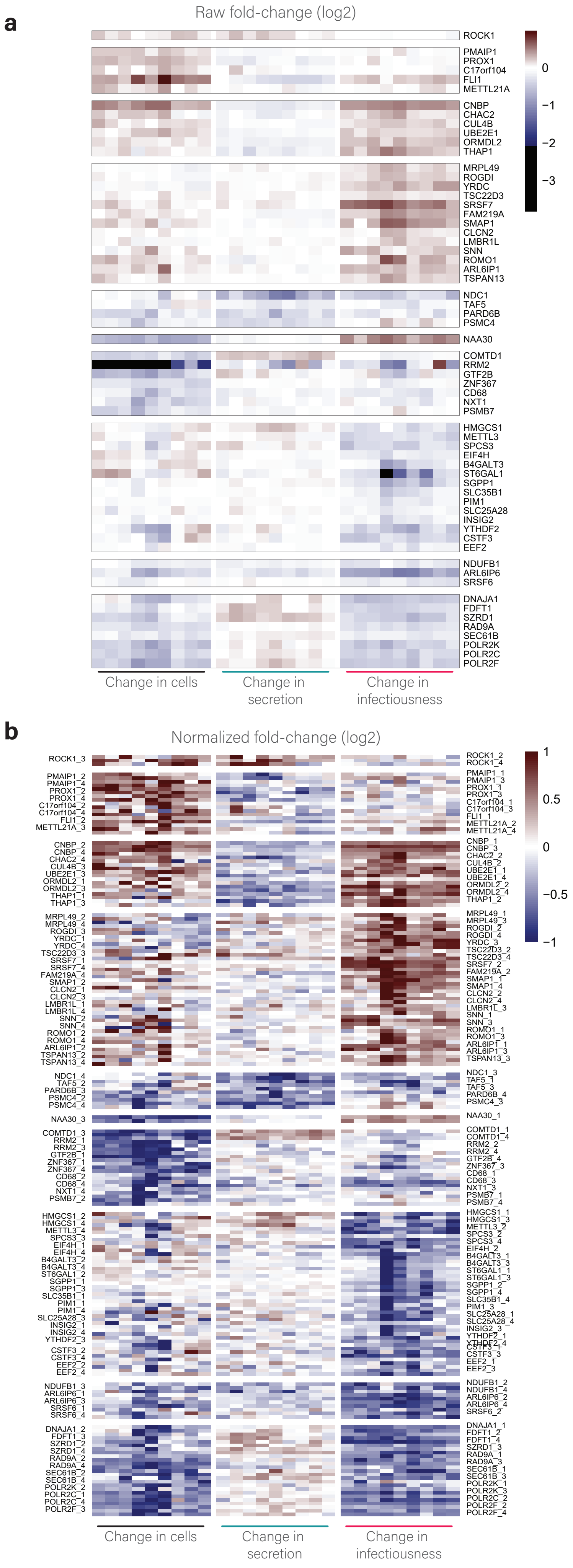

### Supplementary figure 5

#### Supplamentary figure 5

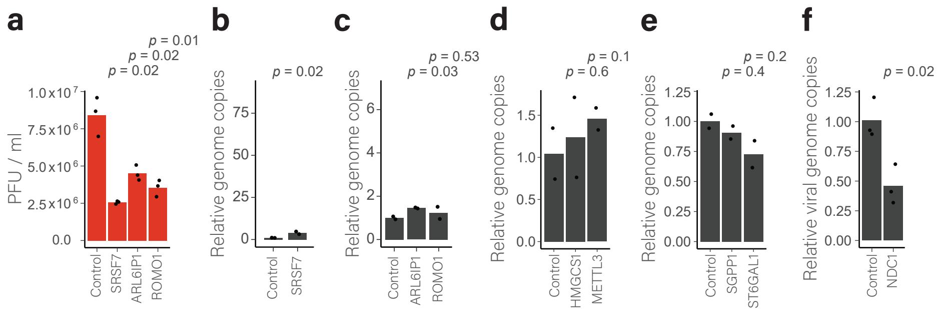
